# Short- and Long-Term Effects of Social Isolation on Adult Murine Bone are Sex-Dependent

**DOI:** 10.64898/2026.04.28.721448

**Authors:** W. Aidan Martel, S. Bradley King, Eleanor Buchanan, Brooklynn M. Merrill, Julia Patrizia Stohn, Daniel J. Brooks, Deborah Barlow, Katherine J. Motyl, Rebecca V. Mountain

## Abstract

Social isolation is a known modifiable risk factor for many chronic diseases including cardiovascular, metabolic, and neurological disorders. Recent research has demonstrated that social isolation is similarly detrimental to skeletal health, but these effects may be sexually dimorphic. In rodents, isolation negatively affects bone in adult male mice, but not in females. However, these sex differences have not been systematically investigated, and it is unknown if they persist with long-term social isolation. The goal of our study was to investigate if isolation-induced bone loss may occur on different timescales between female and male mice, as well as investigate the potential roles of estrogen and testosterone. We examined bone changes in grouped (4 mice/cage) or isolated (1 mouse/cage) female and male 16-week-old C57BL/6J mice after 2, 4, or 8 weeks of treatment. We found that social isolation through single housing significantly reduced bone parameters across treatment lengths in male mice (20% average reduction in Tb.BV/TV; 8% average reduction in Ct.Th.) but not in females even with prolonged isolation. In trabecular bone, isolation affected male bone parameters in as little as 2 weeks. Isolation also decreased biomechanical properties in the femur of male but not female mice. While the females’ overall bone phenotype was unaffected, isolated females did show an increase in bone turnover markers with 2 weeks of isolation. Isolation also altered estrogen-related gene expression in male mice isolated for 4 or 8 weeks. Overall, our results demonstrate that both short- and long-term social isolation has sexually dimorphic effects on murine bone. These findings have important clinical implications for individuals at risk for social isolation, as well as for pre-clinical rodent models utilizing single housing.

**Lay Summary:** Social isolation may have worse effects on bone in males than females, but this has not been thoroughly explored. We investigated if different lengths of isolation had different effects on the bones of male and female mice. We found that both short- and long-term isolation had a rapid and negative impact on the bones of male but not female mice, although isolated females did show some changes in markers of bone formation and resorption. These findings have important implications for humans at risk for social isolation, as well as for studies with rodents that use single housing.

## Introduction

Social isolation is emerging as a key modifiable risk factor globally for many chronic diseases, including cardiovascular^1^ and metabolic diseases^2^, mental health disorders^3^, as well as neurodegenerative diseases^4^. A handful of studies, including our own^5,6^, have demonstrated that social isolation is similarly detrimental to skeletal health in rodent models^7-11^. In our previous study on 16-week-old C57BL/6J mice, we found that four weeks of social isolation significantly reduced trabecular and cortical bone in males^5^. We also found that the effects of isolation on bone cannot be explained by thermal stress from single housing^6^. Other groups have found similarly that in rodent models, social isolation negatively impacts trabecular bone parameters^10^, decreases markers of osteoclast activity^7^, decreases bone mineral content (BMC) and bone mineral density (BMD)^9^, reduces periosteal bone formation and bone mass^8^, and decreases adaptive response to artificial loading^11^. Additionally, our general findings of lower bone mass with social isolation are consistent with other rodent stress models, including chronic subordinate colony housing^12^, early life stress^13^, and chronic mild stress^14^, which also lead to bone loss. Accumulating evidence also suggests social isolation may have similar negative effects on human skeletal health, including increased risk for incident fractures^15,16^, lower BMD^17^, and increased risk of osteoporosis^16-19^ and osteopenia^17,20^.

While several studies now support our finding of a negative effect of isolation on bone parameters, our previous study in mice^5^ found that this effect was sex-specific, with social isolation reducing bone mass in male mice but having no significant effect in female mice. Furthermore, while isolated male mice had reductions in both formation and resorption-related gene expression after four weeks of isolation^5^, females showed increased resorption-related gene expression despite lacking any evidence of bone loss. These data, however, represent only a single time point, and it is possible that the protective effects of estrogen on bone^21^ may cause a delay in isolation-induced bone loss in females relative to males, and that females could experience bone loss with a longer isolation treatment. Both estrogens and androgens have been shown to play a critical role in protecting and maintaining bone health, influencing aspects of metabolism and homeostasis^22^. Estrogen preserves bone density, inhibiting bone resorption via osteoclast activity, and promotes osteoblast activity, while androgens contribute by stimulating bone formation both directly and through its conversion to estrogen by aromatase in bone tissue, to maintain a balance between bone resorption and formation^23^. Only one other rodent study^11^ has considered both males and females, but also found worsened bone parameters in isolated male mice and not isolated females. The authors, however, attributed the male-specific isolation-induced bone loss to lack of fighting and skeletal loading in isolated males. Our prior studies have not supported this hypothesis, as we found no differences in weight, fat, or fat-free mass that could indicate substantial differences in physical activity or loading between the isolated and grouped males^5,6^.

The aim of the present study was to investigate how isolation-induced bone loss may occur on different timescales between male and female mice and test the hypothesis that females have a delay in social isolation-induced bone loss relative to males. We also investigated the potential role of estrogen and testosterone in the effects of isolation on bone. Briefly, we found that social isolation rapidly worsened bone parameters in male mice in as little as 2 weeks. Isolation did not affect the bone phenotype in females even with long-term isolation, but isolation did increase bone turnover markers in females, particularly in short-term social isolation. Isolated male mice also had evidence of altered estrogen signaling. Overall, our findings demonstrate key sex- and time-specific effects of social isolation on bone in mice.

## Methods

### Mouse Model

Eight-to 10-week-old male and female C57BL/6J mice (N=96) were obtained from the Jackson Laboratory (Strain #000664). To reduce any potential stress-related effects from shipment, all mice were acclimated in our facility for 6-8 weeks after arrival and prior to the start of the experiment. Mice were housed in the AAALAC-accredited barrier animal facility at MaineHealth Institute for Research (MHIR) on a 14 h light/10 h dark cycle and provided regular chow (Teklad global 18% protein diet, #2918, Envigo) and water ad libitum. All rodent-related procedures in this study were approved by the MHIR Institutional Animal Care and Use Committee. At 16 weeks of age, male and female mice were randomized into grouped (4 mice/cage) or isolated (1 mouse/cage) housing and one of three treatment lengths: 2 weeks, 4 weeks, or 8 weeks of treatment (N=8/group). Our lab has previously published the micro-CT, CTX-I and P1NP, and a portion of the gene expression data from the 4-week treatment length^5^. Shepherd shacks were provided to both grouped and isolated mice in all treatment lengths as enrichment.

### Bone Parameters

Dual energy X-ray absorptiometry (DXA) was performed at baseline for all mice. Areal total body (post-cranial) and femoral BMD (aBMD) were measured as well as total body fat-free and fat mass using a Hologic Faxitron UltraFocus DXA system. Conducting baseline DXA allowed us to confirm there were no significant mean differences in body composition parameters between experimental groups before the experiment began.

High resolution micro-CT (μCT) (vivaCT 40, Scanco Medical AG) was used to assess trabecular and cortical bone volume, mineral density, and microarchitecture of the femur and L5 *ex vivo*. All bone scans were acquired with an isotropic voxel size of 10.5 μm^3^, 70 kVp peak X-ray tube intensity, a 114 mA X-ray tube current, and 250 ms integration time. After dissection and fixation (formalin for 28 h, transferred to 70% ethanol), cortical and trabecular bone were assessed at the femoral mid-diaphysis and distal femur metaphysis, respectively. The femoral cortical region of interest (ROI) began at 55% of the total bone length distal to the femoral head, extending 525 μm distally, and was segmented with a threshold of 700 mg HA/cm^3^. The femoral trabecular ROI started 210 μm proximal to the break in the distal growth plate and extended 1575 μm proximally and was segmented at 350 mg HA/cm^3^. Four femora were held in a custom-made sample holder for each scan. The L5 ROI started 105 μm below and above the endplates and extended across the entire vertebral body, with a segmentation threshold of 375 mg HA/cm^3^. Vertebral columns were scanned individually. Gaussian filtration and segmentation were performed on all scans, and all analyses were performed using Scanco μCT Evaluation Program V6.6.

Three-point bending was performed to measure the mechanical properties of the femur using an electrical force materials testing machine (Electroforce 3230, Bose Corporation, Eden Prairie, MN). The test was performed with the load point in displacement control moving at a rate of 0.1 mm/sec with force and displacement data collected at 60 Hz. The bending fixture had a bottom span length of 8 mm, and all bones were positioned in the same orientation during testing with the cranial surface resting on the supports and being loaded in tension. Bending rigidity (EI, N-mm^2^), apparent modulus of elasticity (E_app_, MPa), ultimate moment (M_ult_, N-mm), apparent ultimate stress (σ_app_, MPa) work to fracture (W_frac_, mJ), and apparent toughness (U_app_, mJ/mm^3^) were calculated based on the force and displacement data from the tests and the mid-shaft geometry measured with μCT (see above). The minimum moment of inertia (I_min_) from μCT was used when calculating the apparent material properties.

Serum bone turnover markers CTX-I and P1NP concentrations were measured with the RatLaps CTX-I and Rat/Mouse P1NP enzyme immunoassays (EIA, Immunodiagnostic Systems), respectively. Blood collected at the time of decapitation was allowed to clot at room temperature for at least 10 minutes and then centrifuged at 10000 g for 10 minutes. Serum was then collected and stored at -80°C. Assays were performed following the manufacturer’s instructions and read on a FlexStation 3 plate reader (Molecular Devices). Results were determined using the FlexStation software with a 4-parameter logistic curve. Samples with a coefficient of variation exceeding 20% were excluded from analysis resulting in a N=4-8 per group.

Bone turnover- and sex hormone-related gene expression was obtained from extracted RNA from whole bone using real-time qPCR (RT-qPCR) following previously described protocols^5^. Briefly, whole tibia were dissected and cleaned of tissue at the time of sacrifice and flash frozen in liquid nitrogen, then stored at -80°C. Samples were pulverized under liquid nitrogen conditions and homogenized in TriReagent (MRC). Samples were incubated in chloroform and centrifuged to induce phase separation. The aqueous layer was isolated and isopropanol was added before being frozen overnight at -80°C. The following day, samples were centrifuged and the supernatant was removed. The RNA pellet was washed in 75% ethanol twice, and then the pellet was allowed to air dry and dissolved in nuclease-free H_2_O. Samples were again frozen overnight at -80°C, and RNA concentration was determined using the NanoDrop 2000 and diluted if greater than 1000 ng/uL with nuclease-free H_2_O. 1000 ng of RNA was added to each cDNA reaction using a high-capacity cDNA reverse transcription kit (Thermo Fisher Scientific). Reverse transcription was performed using a thermal cycler and diluted with nuclease-free H_2_O. RT-qPCR was performed using 3 µl of cDNA, nuclease-free water, SYBR green (BioRad), appropriate forward and reverse primers, and run on a BioLab Laboratories CFX 384 real-time PCR system. Primers were obtained from either Integrated DNA Technologies or Qiagen, and all primer sequences used are given in **Supplemental Table 1**. Gene expression was calculated using the ΔΔCq method^24^. No housekeeping gene was identified that was consistent across all treatment lengths, therefore *Actb* was used as a housekeeping gene for the 2-week treatment length, and *Hprt* was used as a housekeeping gene for the 4- and 8-week treatment lengths. Any Cq values >2 SD above or below the group mean were excluded as outliers.

### Circulating Sex Hormone Levels

Circulating estradiol in females and testosterone in males were measured in serum using liquid chromatography-tandem mass spectrometry (LC-MS/MS)^25^. Briefly, testosterone and estradiol were simultaneously extracted from mouse serum using a validated extraction method with a range of 0.100-50.0 nM for both analytes. The method employed isotope dilution to correct for any differences in recovery. Supported liquid extraction (SLE) plates were used to emulate traditional liquid/liquid extraction of testosterone and estradiol. Serum was adjusted to pH 5.5 with 500 mM ammonium acetate and applied to the SLE plate and allowed to adsorb. Analytes were eluted with a 1:3 mixture of hexane and ethyl acetate. The eluate was then dried under nitrogen and reconstituted in 100mM sodium bicarbonate. A solution of 10 mg/mL dansyl chloride was then added and the entire reconstituted sample was incubated at 65°C to derivatize estradiol and its deuterated internal standard. Testosterone and its deuterated internal standard do not react with dansyl chloride. The resulting solution was then centrifuged and injected for LC-MS/MS analysis. Separation was achieved via gradient elution on a reversed phase column using mobile phases of 0.2% formic acid in water and methanol with 10 mM ammonium acetate. Detection was accomplished via positive ion multiple reaction monitoring. Calibrants prepared in serum were used to form a calibration curve using 1/x^2^ weighting, plotting peak area ratio of analyte:internal standard. Sample peak area ratios were interpolated from the calibration curve to determine concentration.

### Statistical Analyses

All statistical analyses for rodent data were performed using GraphPad Prism 19 XML statistical software. α≤0.05 was considered statistically significant. Two-way ANOVA was used to test for significant main effects or interactions of housing and/or treatment length (time). Tukey’s post hoc test was performed for multiple comparisons for any data with a significant interaction effect. Multiple t-tests were used to analyze all gene expression results to avoid any batch effects between isolated cDNA.

## Results

### Sex differences in the effects of social isolation on bone persist with 8 weeks of social isolation

Our µCT results showed there was no significant main effect of social isolation on femoral trabecular and cortical bone parameters in female mice (**Figure 1A-B**), apart from a significant increase in cortical porosity with isolation (**Supplemental Table 3**). There was a significant main effect of time, which can be interpreted as the effect of age in the mice, in multiple femur parameters in females, with a decrease in femoral Tb.BV/TV, Tb.BMD, Conn. D, Tb.N. and increase in Tb.Sp., Ct.Ar., Ct.TMD, and pMOI (**Supplemental Tables 2 and 3**). Females showed similar effects in the fifth lumbar vertebrae, with no significant effects of housing or significant interaction effects, but with time (i.e., age) having a significant main effect on Tb.BMD, BS/BV, and Tb.Th.(**Figure 1E-F, Supplemental Table 4**).

**Figure 1.**
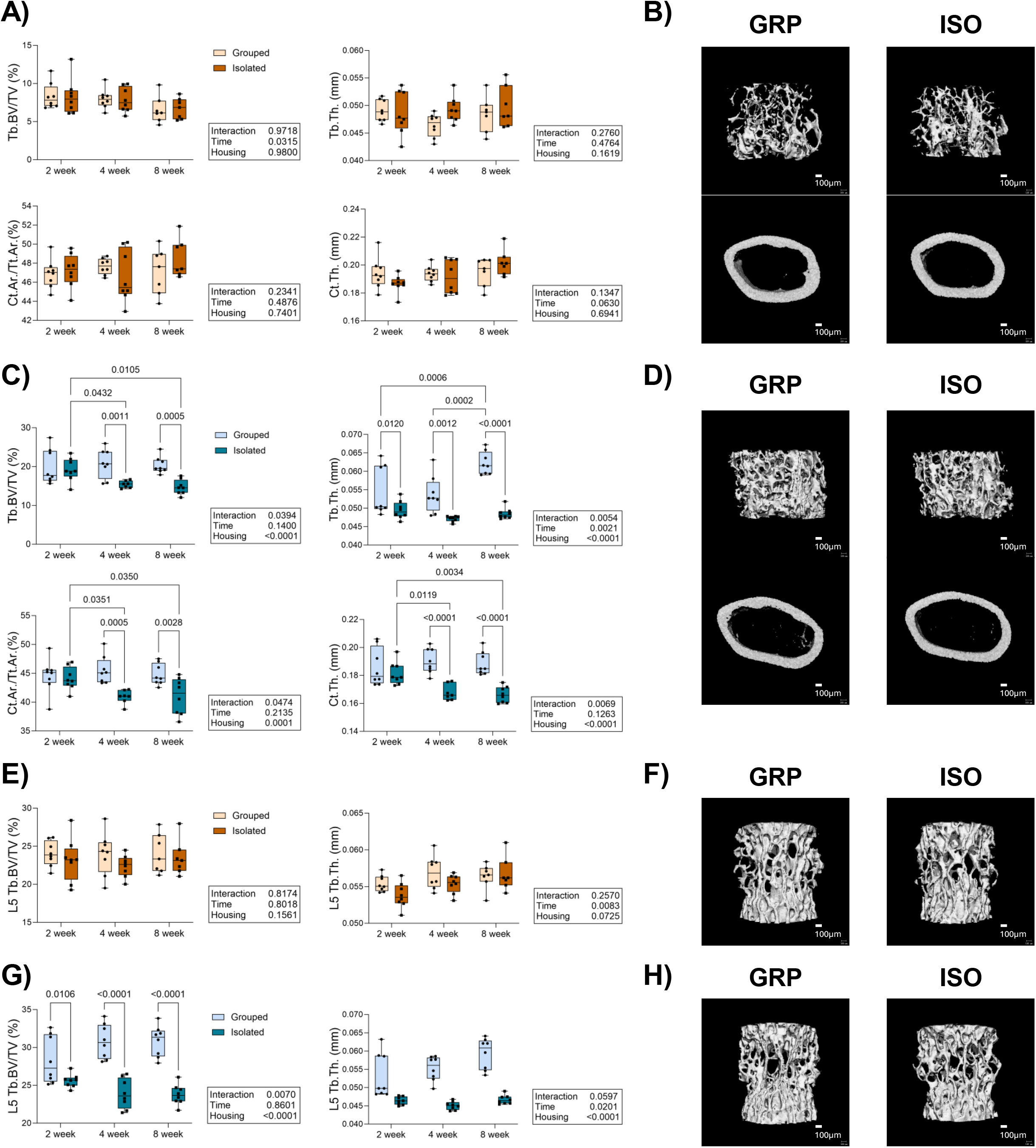
Social isolation decreased trabecular and cortical parameters in the femur and vertebrae of male mice across treatment lengths. **(A-D)** Trabecular and cortical bone parameters were measured in the femur of grouped (GRP) and isolated (ISO) female (A-B) and male (C-D) mice. **(E-H)** Trabecular parameters were measured in the fifth lumbar vertebrae of female (E-F) and male (G-H) mice. All measures were obtained via high resolution µCT (vivaCT 40, Scanco Medical AG). Representative images from 8-week treatment length are shown in panels **B, D, F**, and **H**, scale bar = 100µm. See **Supplemental Tables 2-4** for all µCT parameters. Tb.BV/TV = trabecular bone volume/total volume; Tb.N = trabecular number; Tb.Th = trabecular thickness; Tb.Sp = trabecular separation; Ct. = cortical; Ar. = area; Tt. = total; Ct.Ar/Tt.Ar= cortical area/total area; Ct.Th. = cortical thickness. *P*-values for 2-way ANOVA interaction and main effects presented in boxes to the right of graphs, pairwise comparisons with p < 0.05 shown on graphs for significant interaction effects, N=7-8 group.

Unlike in females, social isolation significantly worsened most femoral parameters in male mice, including Tb.BV/TV, Tb.BMD, Tb.BS/BV, SMI, Tb.Th., Ct.Ar., Ct.Ar/Tt.Ar, and Ct.Th independent of time (**Figure 1C-D, Supplemental Tables 2** and **3**). On average, social isolation decreased femoral Tb.BV/TV and Ct.Th. by 20% and 8% respectively across treatment lengths. There was a significant interaction between housing and time in Tb.BV/TV, Tb.BMD, Tb.BS/BV, Conn.D., SMI, Tb.Th., Ct.Ar/Tt.Ar., and Ct.Th of the femur, with the majority of significant effects with 4 and 8 weeks of isolation. Surprisingly, isolation significantly decreased femoral Tb.Th. and increased Conn.D. in as little as 2 weeks of isolation. Time also had a significant main effect on some trabecular parameters in the male femur, including decreasing Conn.D, Tb.N., and increasing Tb.Th., and Tb.Sp with age. In the male L5 vertebrae, social isolation significantly worsened Tb. BV/TV, Tb.BMD, BS/BV, SMI, Tb.Th, and Tb.Sp, and increased Conn.D (**Figure 1G-H, Supplemental Table 4**). Isolation reduced vertebral Tb.BV/TV by approximately 12% across all treatment lengths. Time and housing had a significant interaction effect in Tb.BV/TV, Tb.BMD, and SMI. Like the femur, isolation significantly decreased male vertebral Tb.BMD and increased SMI in only 2 weeks. Isolation significantly decreased male vertebral Tb.BV/TV with 4 and 8 weeks of treatment. Time also had a significant main effect on Tb.BS/BV, Conn.D., Tb.N, Tb.Th., and Tb.Sp.

We next investigated how social isolation may impact biomechanical properties of the femur. Three-point bending was used to evaluate bone structural and apparent material properties. There was no significant main effect of social isolation on female mechanical parameters or significant interactions, although there was a significant main effect of time on several parameters, with bending rigidity increasing with age, and post-yield displacement, work to fracture, and apparent toughness to fracture worsening with age (**Figure 2A, Supplemental Table 5**). Conversely, social isolation worsened bending rigidity (∼12% reduction), post-yield displacement (∼27% reduction), work to fracture (∼26% reduction), and apparent toughness to fracture (∼23% reduction) in male mice independent of time (**Figure 2B, Supplemental Table 5)**. There was no significant main effect of time or significant interaction effect in the mechanical properties of male femurs.

**Figure 2.**
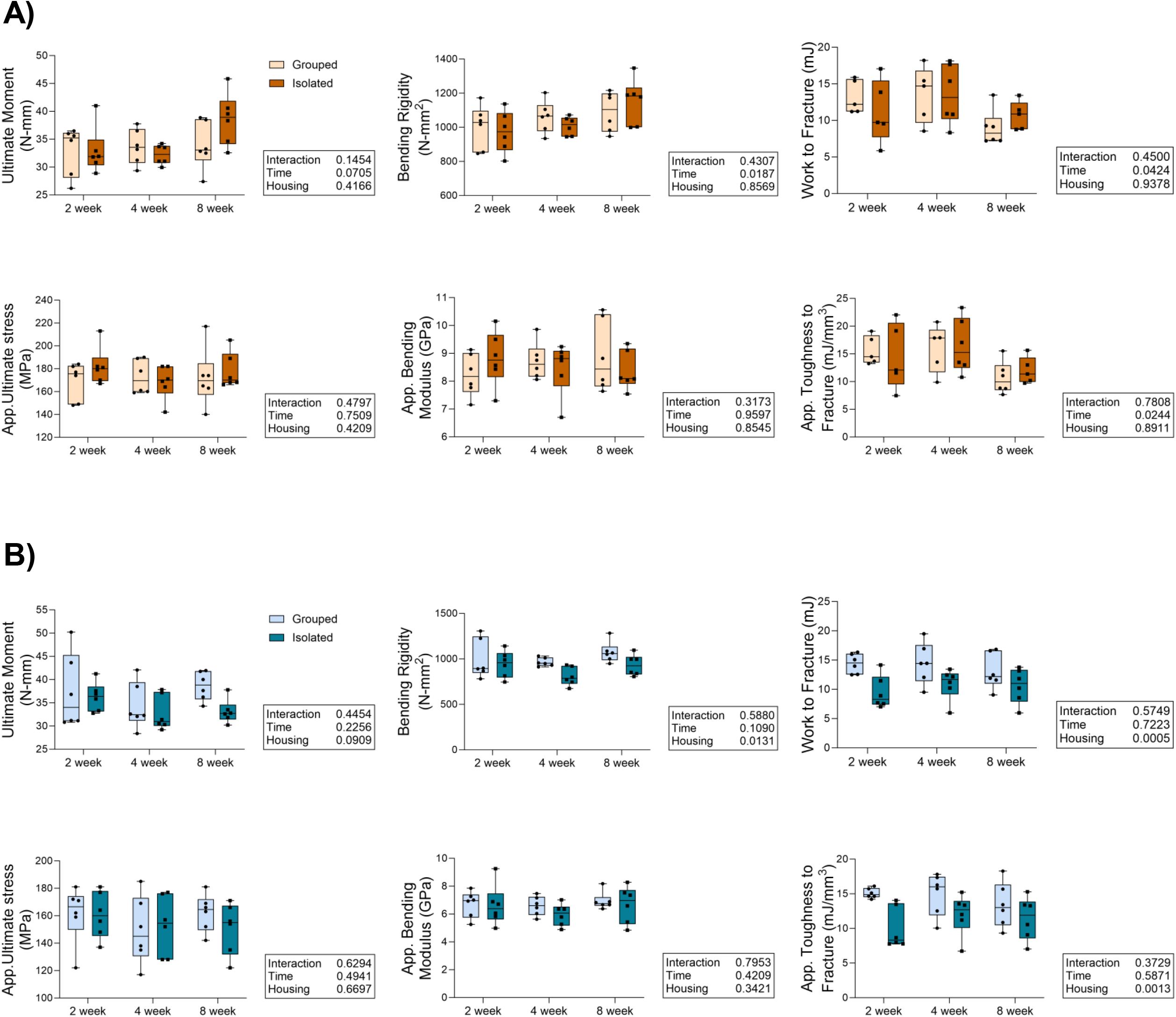
Social isolation decreased femur mechanical properties across treatment lengths in male mice. Femur structural and material properties were measured by three-point bending in females **(A)** and males **(B)**. See **Supplemental Table 5** for all measured parameters. App. = apparant; Disp. = displacement; Ult. = ultimate. N=5-6/group. *P*-values for 2-way ANOVA interaction and main effects presented in boxes to the right of graphs, pairwise comparisons with p < 0.05 shown on graphs for significant interaction effects.

### Social isolation effects on bone turnover markers vary by sex and time

To investigate the mechanisms of social isolation-induced bone loss further, we measured circulating markers of bone turnover in serum, as well as formation- and resorption-related gene expression in whole bone in female and male mice. In females, social isolation significantly increased circulating P1NP, a marker of bone formation (**Figure 3A**). P1NP also increased with time independent of housing conditions in females. Social isolation’s effects on CTX-I, a marker of bone resorption, differed depending on the length of treatment. Isolated females had significantly higher CTX-I levels than grouped females with 2 weeks of treatment, but there were no significant differences between treatments at 4 or 8 weeks. Among the grouped female mice, CTX-I peaked during the 4-week treatment, increasing between the 2- and 4-week treatments, but declining between the 4- and 8-week treatments. Isolated females showed similar levels between the 2- and 4-week treated mice but had a significant decline in the 8-week treated mice relative to both 2- and 4-week mice. There was also a significant main effect of time, with average CTX-I levels declining with age.

**Figure 3.**
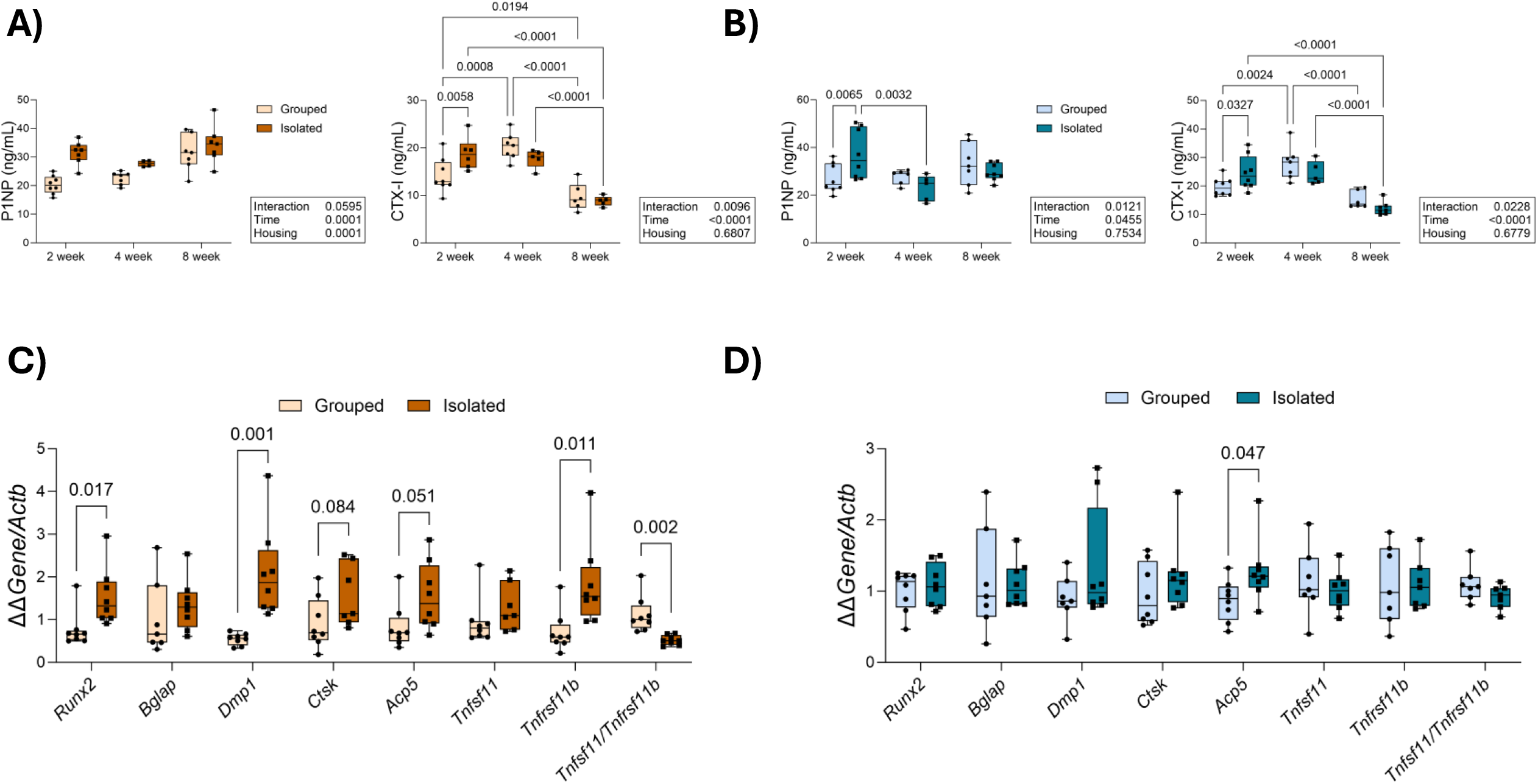
Effects of social isolation on bone turnover markers vary by sex and treatment length. **A-B)** Circulating bone turnover markers measured in serum with enzyme immunoassays. *P*-values for 2-way ANOVA interaction and main effects presented in boxes to the right of graphs, pairwise comparisons with p < 0.05 shown on graphs for significant interaction effects. N=4-8/group. **C-D**) 2-week female (C) and male (D) gene expression, normalized to *Actb*, from whole tibia was measured using RT-qPCR. See **Supplemental Figure 1** for additional gene expression data. Analyzed via multiple t-tests, comparisons with p < 0.1 shown on graphs. N=6-8/group.

In male mice, there was no significant main effect of isolation in either P1NP or CTX-I, but there was a significant interaction between housing and time (**Figure 3B**). Pairwise comparisons demonstrate that social isolation significantly increased P1NP and CTX-I with 2 weeks of treatment, but similar to the female CTX-I, there were no significant differences between treatment at 4 or 8 weeks. Time also had a significant main effect on P1NP levels and declined between 2 and 4 weeks of treatment in the isolated male mice specifically. Like the females, CTX-I significantly increased among the grouped males between 2 and 4 weeks of treatment but significantly declined between 4 and 8 weeks. 8-week isolated males also showed significantly lower CTX-I levels relative to 2- and 4-week treated mice. CTX-I also significantly declined with time regardless of housing condition.

We further investigated changes in bone turnover via gene expression from whole bone. Focusing on changes with 2 weeks of isolation (i.e., gene expression changes preceding the majority of observed bone changes in males), like the P1NP results, social isolation significantly increased bone formation-related gene expression in female mice, increasing expression of *Runx2*, a gene important in osteoblast differentiation, and *Dmp1*, a gene important in osteoblast function and mineralization (**Figure 3C**). Isolation also increased resorption-related gene expression including *Ctsk*, expressed by osteoclasts and important in breaking down bone, and *Acp5*, which encodes TRAP, although this did not reach statistical significance. Isolation significantly increased Osteoprotegerin (OPG) (encoded by *Tnfrsf11b*) and decreased the RANKL/OPG ratio (*Tnfsf11/Tnfrsf11b*), indicating a shift to more bone formation and less resorption with isolation. Similar effects in formation- and resorption-related gene expression were seen at 4 weeks of isolation, but RANKL/OPG was higher in isolated mice relative to grouped mice (**Supplemental Figure 1A**). By 8 weeks of isolation, formation-related expression was still elevated in isolated females, but *Acp5* was significantly lower, and there were no differences in RANKL/OPG (**Supplemental Figure 1C**).

In males, 2 weeks of isolation significantly increased *Acp5* expression, but had no other significant effects on formation- or resorption-related gene expression (**Figure 3D**). However, 4 weeks of isolation significantly decreased *Dmp1, Ctsk*, and *Acp5* expression and decreased *Runx2* expression as well as the RANKL/OPG ratio, although this did not reach statistical significance (**Supplemental Figure 1B**). These data collectively indicate a decrease in bone turnover with 4 weeks of isolation. 8 weeks of isolation significantly increased *Bglap* and *Dmp1* expression and decreased the RANKL/OPG ratio, but as in the 4-week mice, this did not reach statistical significance (**Supplemental Figure 1D**).

### Social isolation alters estrogen receptor expression in males

To determine if the sex differences in isolation-induced bone loss may be a result of differences in sex hormone signaling, we examined circulating levels of estradiol and testosterone in serum, as well as estrogen and androgen receptor gene expression in whole bone. Circulating estradiol levels were not significantly different between isolated and grouped female mice with 4 weeks of treatment (**Figure 4A**), however our sample size was very limited (N=4-5/group). Estradiol was below the quantifiable threshold (<0.100 nM) in male mice as well as in the 2- and 8-week treated female mice. There was no significant main effect of social isolation on circulating testosterone in male mice (**Figure 4B**), but there was a significant main effect of time, with testosterone increasing with age. Testosterone in female mice was below the quantifiable threshold (<0.100 nM) at all time points. Similarly, social isolation had only limited effects on estrogen or androgen receptor expression in females. Social isolation increased expression of estrogen receptor α (ERα) (*Esr1*) and decreased expression of the G protein-coupled estrogen receptor (GPER) (*Gper1)* with 4 weeks of isolation (**Figure 4C**). 2 weeks of isolation also decreased androgen receptor *(Ar*) expression in females, but this effect did not persist with longer treatment lengths (**Supplemental Figure 1E**). In males, however, social isolation significantly reduced the expression of *Esr1* and *Gper1* and tended to increase expression of estrogen receptor β (ERβ) (*Esr2*) with 4 weeks of isolation, although this did not reach statistical significance (**Figure 4D**). *Esr1* expression remained significantly lower in isolated males with 8 weeks of treatment, and *Esr2* and *Gper1* remained trending higher and lower respectively (**Supplemental Figure 1H**). There were no significant differences in androgen receptor expression between isolated or grouped males at any time point.

**Figure 4.**
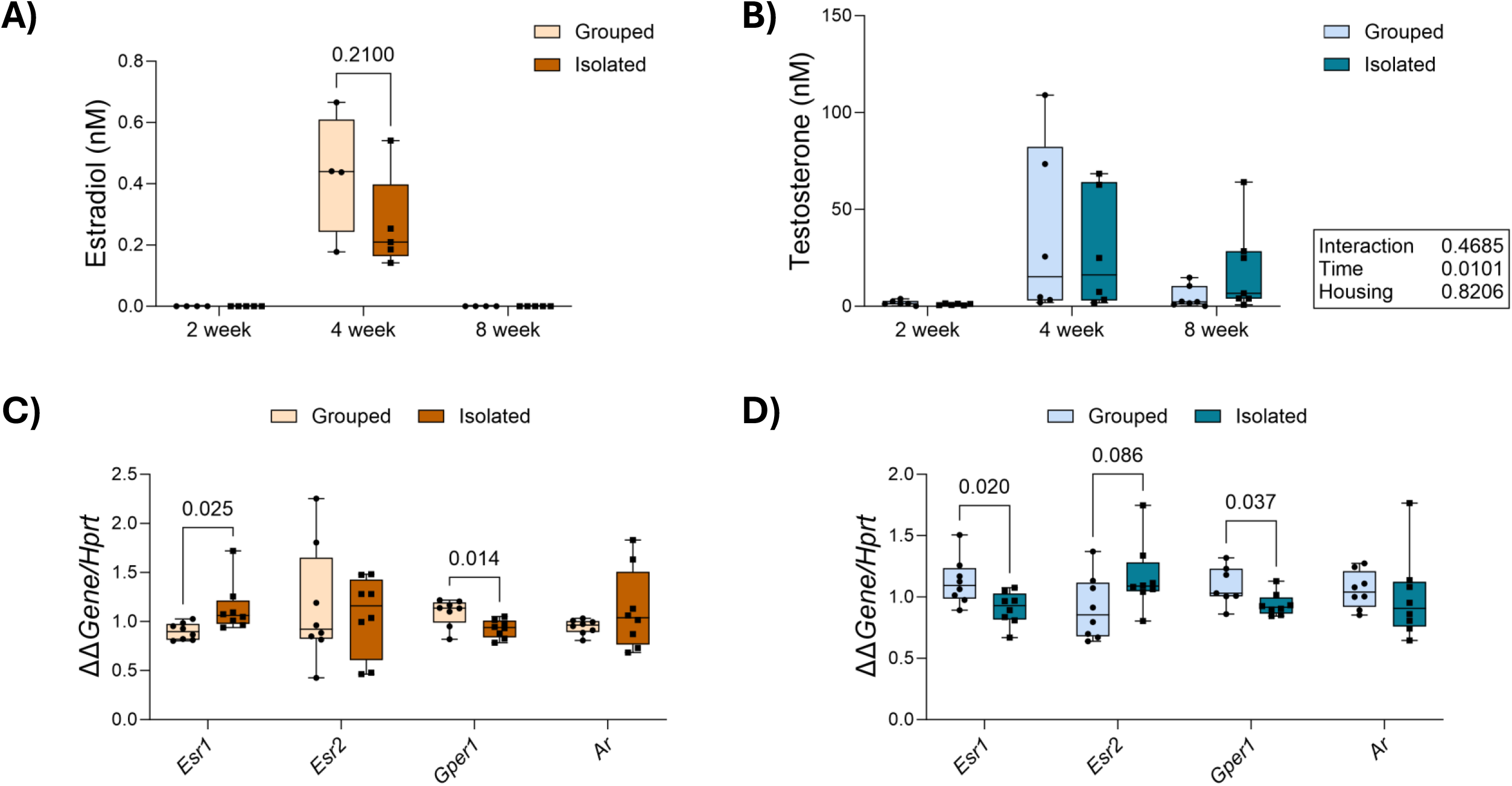
Social isolation did not significantly affect circulating sex hormone levels across treatment lengths but altered estrogen receptor gene expression. **A)** Circulating estradiol measured in female mice serum by liquid chromatography-tandem mass spectrometry (LC-MS/MS). 2- and 8-week estradiol was below quantifiable threshold. Analyzed by t-test. N=4-5/group. **B)** Circulating testosterone measured in male mice serum by LC-MS/MS. Analyzed via 2-way ANOVA. N=6-7/group. **C-D)** 4-week female (C) and male (D) gene expression, normalized *Hprt*, from whole tibia was measured using RT-qPCR. See **Supplemental Figure 1** for all gene expression data. Analyzed via multiple t-tests, comparisons with p < 0.1 shown on graphs. N=7-8/group.

## Discussion

Overall, the results of our study demonstrate that both short- and long-term social isolation negatively affects murine bone in a sex-specific manner. Consistent with our previous studies^5,6^, social isolation negatively affected bone parameters in male mice through 8 weeks of isolation. On the contrary, isolation did not cause negative effects on bone in female mice aside from an increase in cortical porosity but did show evidence of elevated bone turnover occurring with as little as 2 weeks of isolation.

Our results indicate that the negative effects of isolation on bone occur relatively rapidly in male mice and do not recover. In both sexes, we see evidence of changes in bone turnover with 2 weeks of isolation, indicating isolation does have a physiological effect in females, but it is not resulting in a change in bone phenotype. Isolated females show elevated markers of bone formation across treatment lengths, which may partially explain their osteoprotective effects. The increased cortical porosity without other signs of bone loss in isolated females may reflect an overall increase in bone turnover^26,27^, as seen in the serum markers and gene expression, or increased vascularization in the cortical bone^26^, similar to that seen with increasing age. However, cortical porosity is challenging to measure with the current resolution of µCT, and future analysis with nano-CT would be needed to confirm these findings.

By only 2 weeks of isolation, we found significant effects on trabecular bone in the femur and vertebrae male mice. By 4 weeks of isolation, there are significant negative effects across male trabecular and cortical bone parameters. However, there were no significant differences in µCT parameters between 4-week isolated males relative to 8-week isolated males. This suggests that the negative effects of isolation on bone in males happen rapidly and then plateau without causing additional bone loss. The increased expression of *Acp5* and *Dmp1* in the 8-week isolated males may indicate a possible compensation effect, but there is no corresponding increase in P1NP or other bone changes at this timepoint.

Isolation similarly compromised biomechanical properties in males across treatment lengths. Isolation specifically compromised bending rigidity, post-yield displacement, work to fracture, and apparent toughness to fracture in males. These measures indicate that isolation is detrimental to the post-yield displacement and energy absorption capabilities of the male femur, potentially due to isolation-induced changes in the bone’s material properties and organic composition^28^. Such changes in the organic properties of the bone may provide insight into the association between social isolation and increased fracture risk without effects on BMD in humans seen by Bevilacqua et al.^29^ and Lee et al.^15^.

We found that social isolation did not affect circulating testosterone or estradiol measurements in male and female mice respectively, although we were unable to measure estradiol in males or testosterone in females due to low levels. While we did not find an effect of isolation on circulating sex hormone levels, potentially due to our limited sample size, we did observe that social isolation reduced expression of *Esr1* and *Gper1* in whole bone in males. This is particularly interesting as ERα is known to be strongly affected by chronic stress in the brain^30^ as well as being the dominant receptor for estrogen’s effects in bone^31^. It is more highly expressed in osteoblasts, promoting osteoblast differentiation and bone formation^32^, but is also present in osteoclasts and osteocytes, inhibiting bone resorption^33^. GPER is known to mediate more rapid signaling responses across the body^34^, including in stress-related brain regions^34^, as well as in bone^35^. The isolation-induced downregulation of both genes in the whole bone of male mice may indicate that both chronic and acute stress-related pathways are activated with social isolation in males. Although the lack of effect of isolation on bone in females could be due to the general osteoprotective effects of estrogen^21^, the minimal changes in estrogen-related gene expression with isolation across treatment lengths suggests that estrogen alone does not explain the observed sex differences. It is also possible that isolation-induced bone loss may be observed in females with an additional stimulus or stressor, which should be investigated in future studies.

While the mechanisms underlying isolation-induced bone loss—and the observed sex differences—require further investigation, there are several possible areas to explore. Wang et al.^16^ recently found that in humans, systemic inflammation partially mediated the association between social isolation and osteoporosis risk. Although the authors did not specifically investigate differences between women and men, previous studies have shown isolation-induced inflammation disproportionally affects men^36,37^. Glucocorticoid signaling is also activated by social isolation^38^ and has known effects on bone^39^, and glucocorticoid inhibition has been shown to rescue bone loss due in a chronic mild stress model^14^. These bone effects are estrogen-dependent, differing between the sexes, and potentially explaining our observed sex differences^21,40^. Additionally, work on other stress models and bone have found an important role of sympathetic nervous system activity via adrenergic signaling^41,42^. Both catecholamine release and adrenergic receptor expression have been shown to differ between the sexes^43,44^. While few studies have examined the effects of isolation on bone, other rodent studies have explored the effects of chronic social isolation on brain and behavior. These studies have similarly found striking sex differences, including in neurotransmitters^45^, corticosterone^46^, and sleep disturbances^47^, all of which could have downstream effects on bone.

In addition to having important implications for understanding the impact of social isolation stress on human health, our findings have critical importance for pre-clinical rodent models that utilize single housing. Single housing is common for many types of pre-clinical models, including specialty diet^48,49^ and unloading^8,50^ models, both frequently used in bone research. Pre-clinical models that rely on single housing should consider the potential effects of social isolation on bone outcomes, particularly when significant sex differences are observed.

Our study had several major strengths. We leveraged multiple techniques including imaging, mechanical testing, circulating markers, and gene expression to examine the effects of isolation on bone metabolism and differences between sexes. Our findings are also consistent with previous mouse studies^5,11^, as well as some recent human studies^15,19^ indicating sex differences in the effect of isolation on bone. There were also several limitations to this study. First, several of our circulating measures including CTX-I, P1NP, and sex hormones had small sample sizes, in some cases N=4/group. Additionally, testosterone levels and estradiol levels were below the quantifiable threshold in females and males respectively, and estradiol was also not quantifiable at 2 or 8 weeks of isolation in females. These limited sample sizes may affect the analysis and interpretation of these measures.

While several human studies have found an association between social isolation and skeletal health in older adults^15-17,19,20,29^, only two of these studies^15,19^ to date have considered sex differences. Future human studies are needed to ascertain if social isolation is a specific risk factor for men’s skeletal health. To further investigate the sex differences in isolation-induced bone loss in mice and given the changes in estrogen receptor gene expression in isolated males, future work should consider ovariectomy, orchiectomy, and estrogen and testosterone supplementation studies in mice to better understand the potential role of sex hormone signaling. Future research should also investigate potential changes in bone composition, including changes in the collagen matrix that may shed light on the mechanisms underlying isolation-induced bone loss. Lastly, other mechanistic studies, such as those considering the effects of inflammation, glucocorticoids, and adrenergic signaling are needed.

Collectively, the results of this study demonstrate that short- and long-term social isolation has sex-specific effects on murine bone. Isolation rapidly and negatively affects both trabecular and cortical bone in adult male mice and alters circulating bone turnover markers in females without corresponding bone loss. Isolation may also alter estrogen signaling in male mice. These findings have important clinical implications for skeletal health among individuals at risk for social isolation, as well as for pre-clinical rodent models utilizing single housing. Future work will focus on replicating our findings in human populations and exploring potential mechanisms underlying social isolation-induced bone loss.

## Acknowledgments

The authors thank Dr. Evan Buettmann for his assistance in the interpretation of the biomechanical data. The authors also thank the staff of the MaineHealth Institute for Research Animal Facility.

## References

1 Valtorta, N. K., Kanaan, M., Gilbody, S., Ronzi, S. & Hanratty, B. Loneliness and social isolation as risk factors for coronary heart disease and stroke: systematic review and meta-analysis of longitudinal observational studies. Heart 102, 1009–1016 (2016).

2 Rosenkilde, S. et al. Loneliness and the risk of type 2 diabetes. BMJ Open Diabetes Res Care 12 (2024). 10.1136/bmjdrc-2023-003934

3 Donovan, N. J. et al. Loneliness, depression and cognitive function in older U.S. adults. International Journal of Geriatric Psychiatry 32, 564–573 (2017). 10.1002/gps.4495

4 Shen, C. et al. Associations of social isolation and loneliness with later dementia. Neurology 99, e164–e175 (2022).

5 Mountain, R. V. et al. Social isolation through single housing negatively affects trabecular and cortical bone in adult male, but not female, C57BL/6J mice. Bone 172, 116762 (2023).

6 Mountain, R. V. et al. Thermoneutral housing has limited effects on social isolation-induced bone loss in male C57BL/6J mice. JBMR Plus 9 (2025). 10.1093/jbmrpl/ziaf088

7 Schiavone, S. et al. Chronic Psychosocial Stress Impairs Bone Homeostasis: A Study in the Social Isolation Reared Rat. Frontiers in Pharmacology Volume 7-2016 (2016). 10.3389/fphar.2016.00152

8 Morey-Holton, E. R., Halloran, B. P., Garetto, L. P. & Doty, S. B. Animal housing influences the response of bone to spaceflight in juvenile rats. Journal of Applied Physiology 88, 1303–1309 (2000). 10.1152/jappl.2000.88.4.1303

9 Nagy, T. R., Krzywanski, D., Li, J., Meleth, S. & Desmond, R. Effect of Group vs. Single Housing on Phenotypic Variance in C57BL/6J Mice. Obesity Research 10, 412–415 (2002). 10.1038/oby.2002.57

10 Tahimic, C. G. T. et al. Influence of Social Isolation During Prolonged Simulated Weightlessness by Hindlimb Unloading. Frontiers in Physiology Volume 10-2019 (2019). 10.3389/fphys.2019.01147

11 Meakin, L. B. et al. Male mice housed in groups engage in frequent fighting and show a lower response to additional bone loading than females or individually housed males that do not fight. Bone 54, 113–117 (2013).

12 Langgartner, D., Füchsl, A. M., Uschold-Schmidt, N., Slattery, D. A. & Reber, S. O. Chronic Subordinate Colony Housing Paradigm: A Mouse Model to Characterize the Consequences of Insufficient Glucocorticoid Signaling. Frontiers in Psychiatry Volume 6-2015 (2015). 10.3389/fpsyt.2015.00018

13 Wuertz-Kozak, K. et al. Effects of Early Life Stress on Bone Homeostasis in Mice and Humans. International Journal of Molecular Sciences 21, 6634 (2020).

14 Henneicke, H. et al. Chronic mild stress causes bone loss via an osteoblast-specific glucocorticoid-dependent mechanism. Endocrinology 158, 1939–1950 (2017).

15 Lee, A. et al. Associations between Social Isolation Index and changes in grip strength, gait speed, bone mineral density (BMD), and self-reported incident fractures among older adults: Results from the Canadian Longitudinal Study on Aging (CLSA). PLOS ONE 18, e0292788 (2023). 10.1371/journal.pone.0292788

16 Wang, X. et al. Social Isolation and Loneliness Elevating Osteoporosis and Fracture Morbidity, Mediated by Frailty and Systemic Inflammation. Biopsychosocial Science and Medicine, 10.1097 (2025).

17 Xu, Z. et al. The association of loneliness with bone mineral density, osteoporosis, osteopenia, fall, and sarcopenia among older adults: results from Mr. and Ms. Os (Hong Kong) study. Osteoporos Int 36, 2471–2481 (2025). 10.1007/s00198-025-07720-w

18 Christiansen, J. et al. Associations of loneliness and social isolation with physical and mental health among adolescents and young adults. Perspectives in Public Health 141, 226–236 (2021). 10.1177/17579139211016077

19 Zhou, J., Hu, X., Zhou, S., Liu, T. & Chen, Z. Social isolation, loneliness, and genetic susceptibility in relation to the risk of incident osteoporosis: a prospective cohort study based on the UK biobank. Int J Surg (2025). 10.1097/js9.0000000000003467

20 Duckworth, E. et al. What personal factors are associated with osteoporosis, fragility fracture, and osteopenia? A population-level analysis using the United Kingdom Biobank. Bone 190, 117277 (2025).

21 Khosla, S., Oursler, M. J. & Monroe, D. G. Estrogen and the skeleton. Trends in Endocrinology & Metabolism 23, 576–581 (2012).

22 Manolagas, S. C., O’Brien, C. A. & Almeida, M. The role of estrogen and androgen receptors in bone health and disease. Nat Rev Endocrinol 9, 699–712 (2013). 10.1038/nrendo.2013.179

23 Almeida, M. et al. Estrogens and Androgens in Skeletal Physiology and Pathophysiology. Physiol Rev 97, 135–187 (2017). 10.1152/physrev.00033.2015

24 Livak, K. J. & Schmittgen, T. D. Analysis of Relative Gene Expression Data Using Real-Time Quantitative PCR and the 2™ΔΔCT Method. Methods 25, 402–408 (2001). 10.1006/meth.2001.1262

25 Satoh, M. et al. Development and validation of the simultaneous measurement of estrone and 17-β estradiol in serum by LC-MS/MS for clinical laboratory applications. Medical Mass Spectrometry 3, 19–29 (2019).

26 Piemontese, M. et al. Old age causes de novo intracortical bone remodeling and porosity in mice. JCI Insight 2 (2017). 10.1172/jci.insight.93771

27 Zebaze, R. et al. Increased Cortical Porosity and Reduced Trabecular Density Are Not Necessarily Synonymous With Bone Loss and Microstructural Deterioration. JBMR Plus 3, e10078 (2019). 10.1002/jbm4.10078

28 Jepsen, K. J., Silva, M. J., Vashishth, D., Guo, X. E. & Van Der Meulen, M. C. Establishing biomechanical mechanisms in mouse models: practical guidelines for systematically evaluating phenotypic changes in the diaphyses of long bones. Journal of Bone and Mineral Research 30, 951–966 (2015).

29 Bevilacqua, G. et al. The association between social isolation and musculoskeletal health in older community-dwelling adults: findings from the Hertfordshire Cohort Study. Quality of life research 30, 1913–1924 (2021).

30 Handa, R. J., Mani, S. K. & Uht, R. M. Estrogen receptors and the regulation of neural stress responses. Neuroendocrinology 96, 111–118 (2012). 10.1159/000338397

31 Börjesson, A., Lagerquist, M. K., Windahl, S. H. & Ohlsson, C. The role of estrogen receptor α in the regulation of bone and growth plate cartilage. Cellular and Molecular Life Sciences 70, 4023–4037 (2013).

32 Almeida, M. et al. Estrogen receptor-α signaling in osteoblast progenitors stimulates cortical bone accrual. The Journal of clinical investigation 123 (2012).

33 Martin-Millan, M. et al. The estrogen receptor-α in osteoclasts mediates the protective effects of estrogens on cancellous but not cortical bone. Molecular endocrinology 24, 323–334 (2010).

34 Prossnitz, E. R. & Barton, M. The G-protein-coupled estrogen receptor GPER in health and disease. Nature Reviews Endocrinology 7, 715–726 (2011).

35 Chuang, S.-C., Chen, C.-H., Chou, Y.-S.Ho, M.-L. & Chang, J.-K. G protein-coupled estrogen receptor mediates cell proliferation through the cAMP/PKA/CREB pathway in murine bone marrow mesenchymal stem cells. International journal of molecular sciences 21, 6490 (2020).

36 Qi, X., Ng, T. K. S. & Wu, B. Sex differences in the mediating role of chronic inflammation on the association between social isolation and cognitive functioning among older adults in the United States. Psychoneuroendocrinology 149, 106023 (2023).

37 Yang, Y. C., McClintock, M. K., Kozloski, M. & Li, T. Social isolation and adult mortality: the role of chronic inflammation and sex differences. Journal of health and social behavior 54, 183–203 (2013).

38 Cacioppo, J. T., Cacioppo, S., Capitanio, J. P. & Cole, S. W. The neuroendocrinology of social isolation. Annual review of psychology 66, 733 (2015).

39 Canalis, E., Mazziotti, G., Giustina, A. & Bilezikian, J. P. Glucocorticoid-induced osteoporosis: pathophysiology and therapy. Osteoporosis International 18, 1319–1328 (2007).

40 Gohel, A.McCarthy, M.-B. & Gronowicz, G. Estrogen prevents glucocorticoid-induced apoptosis in osteoblasts in vivo and in vitro. Endocrinology 140, 5339–5347 (1999).

41 Haffner-Luntzer, M. et al. Chronic psychosocial stress compromises the immune response and endochondral ossification during bone fracture healing via β-AR signaling. Proceedings of the National Academy of Sciences 116, 8615–8622 (2019).

42 Yirmiya, R. et al. Depression induces bone loss through stimulation of the sympathetic nervous system. Proceedings of the National Academy of Sciences 103, 16876–16881 (2006).

43 Gomes, H. L. et al. Influence of gender and estrous cycle on plasma and renal catecholamine levels in rats. Canadian journal of physiology and pharmacology 90, 75–82 (2012).

44 Riedel, K. et al. Estrogen determines sex differences in adrenergic vessel tone by regulation of endothelial β-adrenoceptor expression. American Journal of Physiology-Heart and Circulatory Physiology 317, H243–H254 (2019).

45 Heng, V., Zigmond, M. & Smeyne, R. J. Neuroanatomical and neurochemical effects of prolonged social isolation in adult mice. Frontiers in Neuroanatomy Volume 17-2023 (2023). 10.3389/fnana.2023.1190291

46 Cepeda, Y. et al. Prolonged social isolation promotes depressive-like behavior in male and female mice, with sex-related differences in the stress response. Biol Sex Differ (2026). 10.1186/s13293-026-00874-0

47 Li, S. et al. Time-Dynamic analysis of sex-specific NREM sleep disturbance induced by social isolation among adolescent mice. Transl Psychiatry (2026). 10.1038/s41398-026-03895-w

48 Moneo, M., Martín Zúñiga, J. & Morón, I. Caloric restriction in grouped rats: aggregate influence on behavioural and hormonal data. Lab Anim 51, 490–497 (2017). 10.1177/0023677216686805

49 Choi, B. S. et al. Housing matters: Experimental variables shaping metabolism in obese mice. Mol Metab 98, 102190 (2025). 10.1016/j.molmet.2025.102190

50 Tahimic, C. G. et al. Influence of social isolation during prolonged simulated weightlessness by hindlimb unloading. Frontiers in physiology 10, 1147 (2019).

