## Supplemental Materials for "Short- and Long-Term Effects of Social Isolation on Adult Murine Bone are Sex-Dependent"

**Supplemental Table 1.** Primer sequences used for mouse qPCR.

| **Gene** | **Abbreviation** | **Forward Primer Sequence** | **Reverse Primer Sequence** | **Manufacturer** |
|---|---|---|---|---|
| Acid phosphatase 5 | *Acp5* | NA | NA | Qiagen (#PPM29328F) |
| Beta actin | *Actb* | NA | NA | Qiagen (#PPM02945B) |
| Osteocalcin | *Bglap* | 5'-ACG GTA TCA CTA TTT AGG ACC TGT-3' | 5'-ACT TTA TTT TGG AGC TGC TGT GAC-3' | IDT[1] |
| Cathepsin K | *Ctsk* | NA | NA | Qiagen (#PPM05123C) |
| Dentin matrix acidic phosphoprotein 1 | *Dmp1* | 5'-TCG CTG AGG TTT TGA CCT TGT-3' | 5'-CTC ACT GTT CGT GGG TGG TG-3' | IDT[2] |
| Estrogen receptor α | *Esr1* | NA | NA | Qiagen (#PPM05075C) |
| Estrogen receptor β | *Esr2* | NA | NA | Qiagen (#PPM05186B) |
| G protein-coupled estrogen receptor | *Gper1* | NA | NA | Qiagen (#PPM30630C) |
| Hypoxanthine guanine phosphoribosyl transferase | *Hprt* | 5′-AAG CCT AAG ATG AGC GCA AG-3’ | 5′-TTA CTA GGC AGA TGG CCA CA-3′ | IDT[3] |
| Runt-related transcription factor 2 | *Runx2* | 5'-GAC AGA AGC TTG ATG ACT CTA AAC C-3' | 5'-TCT GTA ATC TGA CTC TGT CCT TGT G-3' | IDT[1] |
| Osteoprotegerin (OPG) | *Tnfrsf11b* | NA | NA | Qiagen (#PPM03404F) |
| Receptor activator of nuclear factor kappa-B ligand (RANKL) | *Tnfsf11* | NA | NA | Qiagen (#PPM03047F) |

1. Ontiveros, C., Irwin, R., Wiseman, R. W. & McCabe, L. R. Hypoxia suppresses runx2 independent of modeled microgravity. Journal of cellular physiology 200, 169-176 (2004).
2. Motyl, K. J. et al. Propranolol attenuates risperidone-induced trabecular bone loss in female mice. Endocrinology 156, 2374-2383 (2015).
3. Vengellur, A. & LaPres, J. The role of hypoxia inducible factor 1α in cobalt chloride induced cell death in mouse embryonic fibroblasts. Toxicological Sciences 82, 638-646 (2004).

**Supplemental Table 2.** Trabecular microarchitecture of distal femur of female and male mice housed in grouped or isolated housing for two, four, or eight weeks. (N=7-8/group).

| | 2 weeks | | 4 weeks | | 8 weeks | | Two-Way ANOVA Results | | |
|---|---|---|---|---|---|---|---|---|---|
| **Trabecular Bone** | **Grouped** | **Isolated** | **Grouped** | **Isolated** | **Grouped** | **Isolated** | **Interaction** | **Time** | **Housing** |
| **Females** | | | | | | | | | |
| Tb.BV/TV(%) | 8.32 ± 1.70 | 8.26 ± 2.30 | 7.98 ± 1.28 | 7.84 ± 1.61 | 6.56 ± 1.74 | 6.72 ± 1.36 | 0.9718 | **0.0315** | 0.9800 |
| Tb.BMD ($mg/cm^3$) | 122.95 ± 13.39 | 123.41 ± 17.01 | 118.33 ± 8.62 | 118.66 ± 12.38 | 109.36 ± 15.99 | 110.85 ± 12.00 | 0.9918 | **0.0376** | 0.8498 |
| BS/BV ($mm^2/mm^3$) | 58.14 ±3.16 | 59.87 ± 6.23 | 60.48 ± 3.97 | 57.75 ± 3.63 | 60.10 ± 5.25 | 58.63 ± 5.29 | 0.3922 | 0.9773 | 0.5545 |
| Conn.D ($1/mm^3$) | 53.19 ± 17.82 | 54.36 ± 20.67 | 60.01 ± 13.02 | 49.26 ± 17.91 | 38.13 ± 14.04 | 35.09 ± 12.59 | 0.5863 | **0.0071** | 0.3904 |
| SMI | 2.94 ± 0.26 | 2.96 ± 0.24 | 2.80 ± 0.17 | 2.94 ± 0.32 | 3.00 ± 0.23 | 3.02 ± 0.25 | 0.7126 | 0.3107 | 0.4244 |
| Tb.N (1/mm) | 3.67 ± 0.26 | 3.79 ± 0.33 | 3.55 ± 0.21 | 3.49 ± 0.18 | 3.20 ± 0.26 | 3.20 ± 0.09 | 0.563 | **<0.0001** | 0.7368 |
| Tb.Th (mm) | 0.049 ± 0.002 | 0.49 ± 0.004 | 0.046 ± 0.002 | 0.049 ± 0.002 | 0.048 ± 0.003 | 0.050 ± 0.004 | 0.276 | 0.4764 | 0.1619 |
| Tb.Sp (mm) | 0.274 ± 0.019 | 0.264 ± 0.019 | 0.281 ± 0.017 | 0.288 ± 0.016 | 0.314 ± 0.023 | 0.313 ± 0.009 | 0.4156 | **<0.0001** | 0.8463 |
| **Males** | | | | | | | | | |
| Tb.BV/TV(%) | 19.92 ± 4.46 | 19.22 ± 2.94 | 20.80 ± 3.72 | 15.05 ± 1.31^†† | 20.32 ± 2.11 | 14.69 ± 1.91^††† | **0.0394** | 0.14 | **<0.0001** |
| Tb.BMD ($mg/cm^3$) | 213.10 ± 36.33 | 204.54 ± 20.88 | 216.18 ± 30.04 | 172.70 ± 11.31^††† | 220.74 ± 17.97 | 170.81 ± 13.91^††† | **0.0361** | 0.1721 | **<0.0001** |
| BS/BV ($mm^2/mm^3$) | 46.99 ± 6.24 | 50.39 ± 3.64 | 46.29 ± 5.13 | 54.44 ± 1.49††† | 40.81 ± 2.90^^ | 52.90 ± 2.48†††† | **0.0137** | 0.0551 | **<0.0001** |
| Conn.D ($1/mm^3$) | 158.49 ± 16.31 | 184.97 ± 25.93†† | 163.49 ± 16.37 | 144.86 ± 19.54^^^ | 112.37 ± 12.24^^^^ | 126.37 ± 22.95^^^^ | **0.0062** | **<0.0001** | 0.2012 |
| SMI | 1.75 ± 0.36 | 1.67 ± 0.35 | 1.53 ± 0.37 | 2.08 ± 0.13^††† | 1.74 ± 0.27 | 2.08 ± 0.26^† | **0.0153** | 0.1885 | **0.0033** |
| Tb.N (1/mm) | 5.33 ± 0.27 | 5.46 ± 0.27 | 5.32 ± 0.26 | 4.96 ± 0.25 | 4.88 ± 0.33 | 4.76 ± 0.30 | 0.066 | **<0.0001** | 0.1534 |
| Tb.Th (mm) | 0.054 ± 0.007 | 0.049 ± 0.002† | 0.054 ± 0.005 | 0.047 ± 0.001†† | 0.062 ± 0.003^^^ | 0.049 ± 0.002†††† | **0.0054** | **0.0021** | **<0.0001** |
| Tb.Sp (mm) | 0.181 ± 0.009 | 0.175 ± 0.009 | 0.180 ± 0.011 | 0.193 ± 0.012 | 0.200 ± 0.013 | 0.203 ± 0.013 | 0.0677 | **<0.0001** | 0.3079 |

Data presented as mean ± SD.

Tb.BV/TV = trabecular bone volume/total volume; Tb.BMD = trabecular bone mineral density; BS/BV = bone surface/bone volume; Conn.D = connectivity density; SMI = structure model index; Tb.N = trabecular number; Tb.Th = trabecular thickness; Tb.Sp = trabecular separation.

Two-way ANOVA: Bold values indicate significance with $p < 0.05$

*Pairwise comparisons for significant interactions:* ^$p < 0.05$, ^^$p < 0.01$, ^^^$p < 0.001$, and ^^^^$p < 0.0001$ compared to housing matched 2-week treated mice.

†$p < 0.05$, ††$p < 0.01$, †††$p < 0.001$, and ††††$p < 0.0001$ compared to time matched grouped mice.

**Supplemental Table 3.** Cortical microarchitecture of femoral midshaft of female and male mice housed in grouped or isolated housing for two, four, or eight weeks. (N=7-8/group).

| | 2 weeks | | 4 weeks | | 8 weeks | | Two-Way ANOVA Results | | |
|---|---|---|---|---|---|---|---|---|---|
| **Cortical Bone** | **Grouped** | **Isolated** | **Grouped** | **Isolated** | **Grouped** | **Isolated** | **Interaction** | **Time** | **Housing** |
| **Females** | | | | | | | | | |
| Ct.Ar ($mm^2$) | 0.823 ± 0.046 | 0.797 ± 0.045 | 0.826 ± 0.027 | 0.828 ± 0.037 | 0.841 ± 0.036 | 0.881 ± 0.051 | 0.1017 | **0.0051** | 0.6615 |
| Ma.Ar ($mm^2$) | 0.931 ± 0.029 | 0.891 ± 0.050 | 0.907 ± 0.040 | 0.954 ± 0.102 | 0.946 ± 0.095 | 0.936 ± 0.071 | 0.2167 | 0.4845 | 0.9626 |
| Tt.Ar ($mm^2$) | 1.753 ± 0.058 | 1.688 ± 0.073 | 1.733 ± 0.061 | 1.781 ± 0.116 | 1.787 ± 0.113 | 1.817 ± 0.098 | 0.1698 | 0.0538 | 0.8682 |
| Ct.Ar/Tt.Ar (%) | 46.893 ± 1.503 | 47.210 ± 1.761 | 47.669 ± 0.836 | 46.569 ± 2.713 | 47.159 ± 2.408 | 48.520 ± 1.996 | 0.2341 | 0.4876 | 0.7401 |
| Ct.Th (mm) | 0.194 ± 0.011 | 0.187 ± 0.006 | 0.194 ± 0.006 | 0.192 ± 0.012 | 0.195 ± 0.010 | 0.202 ± 0.009 | 0.1347 | 0.063 | 0.6941 |
| Ct. TMD ($mg/cm^3$) | 1291.226 ± 12.908 | 1280.773 ± 13.725 | 1295.408 ± 15.290 | 1291.215 ± 13.392 | 1300.471 ± 9.433 | 1302.234 ± 11.192 | 0.4397 | **0.0092** | 0.2668 |
| Ct.Por (%) | 0.415 ± 0.046 | 0.482 ± 0.059 | 0.415 ± 0.030 | 0.442 ± 0.028 | 0.404 ± 0.032 | 0.417 ± 0.059 | 0.2195 | 0.0743 | **0.0095** |
| pMOI ($mm^4$) | 0.374 ± 0.034 | 0.352 ± 0.037 | 0.369 ± 0.026 | 0.389 ± 0.046 | 0.389 ± 0.039 | 0.410 ± 0.040 | 0.2058 | **0.0391** | 0.5568 |
| **Males** | | | | | | | | | |
| Ct.Ar ($mm^2$) | 0.891 ± 0.114 | 0.882 ± 0.063 | 0.894 ± 0.057 | 0.820 ± 0.029 | 0.949 ± 0.051 | 0.835 ± 0.048 | 0.1013 | 0.9669 | **0.0012** |
| Ma.Ar ($mm^2$) | 1.110 ± 0.161 | 1.122 ± 0.126 | 1.188 ± 0.112 | 1.184 ± 0.113 | 1.172 ± 0.104 | 1.217 ± 0.220 | 0.684 | 0.3044 | 0.2205 |
| Tt.Ar ($mm^2$) | 2.002 ± 0.242 | 2.004 ± 0.175 | 2.083 ± 0.164 | 2.004 ± 0.138 | 2.121 ± 0.140 | 2.052 ± 0.264 | 0.8161 | 0.4623 | 0.7669 |
| Ct.Ar/Tt.Ar (%) | 44.586 ± 2.965 | 44.076 ± 1.980 | 42.991 ± 1.145 | 41.003 ± 1.667^††† | 44.786 ± 1.842 | 41.017 ± 3.141^†† | **0.0474** | 0.2135 | **0.0001** |
| Ct.Th (mm) | 0.186 ± 0.013 | 0.181 ± 0.008 | 0.180 ± 0.008 | 0.164 ± 0.004^†††† | 0.188 ± 0.008 | 0.166 ± 0.006^^†††† | **0.0069** | 0.1263 | **<0.0001** |
| Ct. TMD ($mg/cm^3$) | 1255.842 ± 11.539 | 1261.041 ± 14.542 | 1246.138 ± 4.044 | 1239.419 ± 15.496 | 1265.993 ± 14.714 | 1256.412 ± 17.243 | 0.0755 | 0.7147 | 0.0763 |
| Ct.Por (%) | 0.454 ± 0.041 | 0.423 ± 0.055 | 0.441 ± 0.045 | 0.532 ± 0.102 | 0.445 ± 0.072 | 0.489 ± 0.084 | 0.1133 | 0.3698 | 0.1822 |
| pMOI ($mm^4$) | 0.480 ± 0.106 | 0.474 ± 0.066 | 0.504 ± 0.069 | 0.458 ± 0.047 | 0.538 ± 0.060 | 0.478 ± 0.096 | 0.6519 | 0.5528 | 0.1998 |

Data presented as mean ± SD.

Ct. = cortical; Ar. = area; Tt. = total; Ma. = medullary; Ct.Ar/Tt.Ar= cortical area/total area; Ct.TMD = cortical tissue mineral density; Ct.Por. = cortical porosity; Ct.Th. = cortical thickness; pMOI = polar moment of inertia.

Two-way ANOVA: Bold values indicate significance with $p < 0.05$

*Pairwise comparisons for significant interactions:* ^$p < 0.05$, ^^$p < 0.01$, ^^^$p < 0.001$, and ^^^^$p < 0.0001$ compared to housing matched 2-week treated mice.

†$p < 0.05$, ††$p < 0.01$, †††$p < 0.001$, and ††††$p < 0.0001$ compared to time matched grouped mice.

**Supplemental Table 4.** Trabecular microarchitecture of L5 vertebrae of female and male mice housed in grouped or isolated housing for two, four, or eight weeks. (N=7-8/group).

| | 2 weeks | | 4 weeks | | 8 weeks | | Two-Way ANOVA Results | | |
|---|---|---|---|---|---|---|---|---|---|
| **Trabecular Bone** | **Grouped** | **Isolated** | **Grouped** | **Isolated** | **Grouped** | **Isolated** | **Interaction** | **Time** | **Housing** |
| **Females** | | | | | | | | | |
| Tb.BV/TV(%) | 24.014 ± 1.656 | 23.175 ± 2.862 | 23.985 ± 2.789 | 22.395 ± 1.425 | 24.00 ± 2.577 | 23.45 ± 2.294 | 0.8174 | 0.8018 | 0.1561 |
| Tb.BMD (mg/cm$^3$) | 232.468 ± 13.372 | 224.060 ± 23.318 | 257.231 ± 23.931 | 244.352 ± 11.155 | 232.17 ± 21.005 | 229.37 ± 19.195 | 0.7749 | **0.0036** | 0.1654 |
| BS/BV (mm$^2$/mm$^3$) | 39.724 ± 1.190 | 41.075 ± 1.204 | 39.007 ± 1.751 | 40.182 ± 0.891 | 39.05 ± 1.477 | 38.93 ± 1.741 | 0.3116 | **0.0300** | 0.0592 |
| Conn.D (1/mm$^3$) | 142.039 ± 12.819 | 149.774 ± 20.103 | 139.849 ± 849 | 131.200 ± 19.877 | 135.58 ± 21.469 | 125.24 ± 18.860 | 0.4025 | 0.1155 | 0.5372 |
| SMI | 0.480 ± 0.147 | 0.564 ± 0.281 | 0.575 ± 0.218 | 0.665 ± 0.159 | 0.45 ± 0.232 | 0.49 ± 0.219 | 0.9561 | 0.1533 | 0.2488 |
| Tb.N (1/mm) | 4.075 ± 0.202 | 4.129 ± 0.407 | 4.098 ± 0.351 | 3.909 ± 0.225 | 3.99 ± 0.289 | 3.92 ± 0.161 | 0.4967 | 0.3709 | 0.4094 |
| Tb.Th (mm) | 0.055 ± 0.001 | 0.054 ± 0.002 | 0.057 ± 0.002 | 0.055 ± 0.001 | 0.06 ± 0.002 | 0.06 ± 0.002 | 0.2570 | **0.0083** | 0.0725 |
| Tb.Sp (mm) | 0.246 ± 0.013 | 0.241 ± 0.027 | 0.247 ± 0.022 | 0.261 ± 0.016 | 0.25 ± 0.019 | 0.25 ± 0.011 | 0.3922 | 0.2659 | 0.4509 |
| **Males** | | | | | | | | | |
| Tb.BV/TV(%) | 28.360 ± 3.137 | 25.601 ± 0.875† | 29.490 ± 2.527 | 23.848 ± 2.004†††† | 30.83 ± 2.026 | 23.76 ± 1.339†††† | **0.007** | 0.8601 | **<0.0001** |
| Tb.BMD (mg/cm$^3$) | 265.873 ± 27.239 | 239.779 ± 7.982†† | 299.167 ± 23.600 | 250.076 ± 15.226†††† | 290.86 ± 19.441^ | 226.45 ± 10.247†††† | **0.0075** | 0.6484 | **<0.0001** |
| BS/BV (mm$^2$/mm$^3$) | 41.844 ± 3.982 | 47.639 ± 1.305 | 41.311 ± 2.579 | 48.362 ± 1.791 | 37.68 ± 2.343 | 47.07 ± 1.468 | 0.0536 | **0.0057** | **<0.0001** |
| Conn.D (1/mm$^3$) | 218.415 ± 18.486 | 273.003 ± 29.026 | 248.713 ± 22.270 | 259.735 ± 27.085 | 189.81 ± 12.120 | 213.47 ± 21.744 | 0.3247 | **<0.0001** | **<0.0001** |
| SMI | 0.328 ± 0.184 | 0.503 ± 0.091† | 0.349 ± 0.199 | 0.736 ± 0.153^†††† | 0.22 ± 0.118 | 0.61 ± 0.116†††† | **0.0143** | 0.6956 | **<0.0001** |
| Tb.N (1/mm) | 5.347 ± 0.251 | 5.473 ± 0.206 | 5.638 ± 0.330 | 5.359 ± 0.340 | 5.27 ± 0.204 | 5.08 ± 0.251 | 0.0835 | **<0.0001** | 0.1634 |
| Tb.Th (mm) | 0.053 ± 0.006 | 0.046 ± 0.001 | 0.054 ± 0.003 | 0.046 ± 0.001 | 0.06 ± 0.004 | 0.05 ± 0.001 | 0.0597 | **0.0201** | **<0.0001** |
| Tb.Sp (mm) | 0.178 ± 0.011 | 0.175 ± 0.007 | 0.170 ± 0.011 | 0.181 ± 0.012 | 0.18 ± 0.009 | 0.19 ± 0.010 | 0.0695 | **0.0005** | **0.0381** |

Data presented as mean ± SD.

Tb.BV/TV = trabecular bone volume/total volume; Tb.BMD = trabecular bone mineral density; BS/BV = bone surface/bone volume; Conn.D = connectivity density; SMI = structure model index; Tb.N = trabecular number; Tb.Th = trabecular thickness; Tb.Sp = trabecular separation.

Two-way ANOVA: Bold values indicate significance with $p < 0.05$

*Pairwise comparisons for significant interactions:* ^$p < 0.05$, ^^$p < 0.01$, ^^^$p < 0.001$, and ^^^^$p < 0.0001$ compared to housing matched 2-week treated mice.

†$p < 0.05$, ††$p < 0.01$, †††$p < 0.001$, and ††††$p < 0.0001$ compared to time matched grouped mice.

**Supplemental Table 5.** Mechanical properties of femur of female and male mice housed in grouped or isolated housing for two, four, or eight weeks. (N=5-6/group).

| | 2 weeks | | 4 weeks | | 8 weeks | | Two-Way ANOVA Results | | |
|---|---|---|---|---|---|---|---|---|---|
| | **Grouped** | **Isolated** | **Grouped** | **Isolated** | **Grouped** | **Isolated** | **Interaction** | **Time** | **Housing** |
| **Females** | | | | | | | | | |
| Bending Rigidity (N-mm$^2$) | 1000 ± 128 | 973 ± 121 | 1061 ± 93 | 1007 ± 55 | 1091 ± 117 | 1152 ± 132 | 0.4307 | **0.0187** | 0.8569 |
| App. Bending Modulus (GPa) | 8.23 ± 0.76 | 8.82 ± 0.98 | 8.71 ± 0.65 | 8.47 ± 0.93 | 8.88 ± 1.29 | 8.37 ± 0.70 | 0.3173 | 0.9597 | 0.8545 |
| Ultimate Moment (N-mm) | 32.97 ± 4.37 | 32.90 ± 4.20 | 33.66 ± 3.15 | 32.24 ± 1.78 | 33.90 ± 4.27 | 38.58 ± 4.68 | 0.1454 | 0.0705 | 0.4166 |
| App.Ultimate stress (MPa) | 169 ± 16 | 182 ± 16 | 173 ± 15 | 168 ± 15 | 172 ± 25 | 178 ± 16 | 0.4797 | 0.7509 | 0.4209 |
| Yield Load (N) | 12.5 ± 1.8 | 12.5 ± 1.8 | 13.3 ± 2.6 | 12.3 ± 1.8 | 12.9 ± 1.3 | 13.9 ± 2.0 | 0.4496 | 0.4992 | 0.9931 |
| Post-yield Disp (mm) | 0.99 ± 0.21 | 0.98 ± 0.67 | 1.13 ± 0.50 | 1.23 ± 0.58 | 0.62 ± 0.28 | 0.66 ± 0.20 | 0.957 | **0.0283** | 0.7779 |
| Work to Ult. moment (mJ) | 3.17 ± 0.86 | 3.29 ± 0.35 | 3.31 ± 0.51 | 3.13 ± 0.62 | 2.86 ± 0.60 | 3.92 ± 1.47 | 0.1667 | 0.8511 | 0.2321 |
| Work to Fracture (mJ) | 13.19 ± 2.29 | 11.21 ± 4.32 | 13.54 ± 3.84 | 13.53 ± 4.07 | 8.96 ± 2.41 | 10.67 ± 1.95 | 0.45 | **0.0424** | 0.9378 |
| App. Toughness to Ult. Moment (mJ/mm$^3$) | 3.80 ± 0.95 | 4.19 ± 0.45 | 4.01 ± 0.77 | 3.79 ± 0.59 | 3.39 ± 0.71 | 4.25 ± 1.33 | 0.3072 | 0.8846 | 0.2324 |
| App. Toughness to Fracture (mJ/mm$^3$) | 15.61 ± 2.59 | 14.46 ± 5.97 | 15.99 ± 4.28 | 16.42 ± 4.90 | 10.65 ± 2.90 | 11.98 ± 2.41 | 0.7808 | **0.0244** | 0.8911 |
| **Males** | | | | | | | | | |
| Bending Rigidity (N-mm$^2$) | 995 ± 216 | 942 ± 146 | 965 ± 47 | 807 ± 102 | 1072 ± 115 | 931 ± 116 | 0.5880 | 0.1090 | **0.0131** |
| App. Bending Modulus (GPa) | 6.70 ± 0.95 | 6.62 ± 1.46 | 6.58 ± 0.69 | 5.94 ± 0.79 | 6.94 ± 0.63 | 6.67 ± 1.32 | 0.7953 | 0.4209 | 0.3421 |
| Ultimate Moment (N-mm) | 37.30 ± 8.06 | 36.26 ± 3.11 | 34.41 ± 5.00 | 32.75 ± 3.76 | 38.65 ± 3.10 | 33.13 ± 2.55 | 0.4454 | 0.2256 | 0.0909 |
| App.Ultimate stress (MPa) | 161 ± 21 | 160 ± 17 | 149 ± 25 | 153 ± 22 | 162 ± 14 | 151 ± 19 | 0.6294 | 0.4941 | 0.6697 |
| Yield Load (N) | 12.3 ± 1.8 | 12.8 ± 1.9 | 11.4 ± 0.9 | 11.9 ± 2.3 | 13.4 ± 2.7 | 12.1 ± 2.4 | 0.4844 | 0.3843 | 0.8426 |
| Post-yield Disp (mm) | 1.00 ± 0.19 | 0.55 ± 0.21 | 1.13 ± 0.37 | 0.86 ± 0.34 | 0.83 ± 0.30 | 0.75 ± 0.34 | 0.3155 | 0.1535 | **0.0114** |
| Work to Ult. moment (mJ) | 4.13 ± 1.46 | 3.66 ± 0.43 | 4.12 ± 0.92 | 3.64 ± 1.06 | 3.97 ± 1.14 | 3.78 ± 0.97 | 0.9263 | 0.9988 | 0.2801 |
| Work to Fracture (mJ) | 14.41 ± 1.68 | 9.46 ± 2.80 | 14.47 ± 3.52 | 10.90 ± 2.66 | 13.07 ± 3.05 | 10.58 ± 2.96 | 0.5749 | 0.7223 | **0.0005** |
| App. Toughness to Ult. Moment (mJ/mm$^3$) | 4.29 ± 1.28 | 3.82 ± 0.26 | 4.30 ± 0.99 | 4.04 ± 1.29 | 4.08 ± 1.27 | 4.10 ± 1.00 | 0.8546 | 0.9628 | 0.5181 |
| App. Toughness to Fracture (mJ/mm$^3$) | 15.07 ± 0.72 | 9.94 ± 2.98 | 14.95 ± 3.04 | 12.01 ± 2.93 | 13.36 ± 3.24 | 11.44 ± 3.09 | 0.3729 | 0.5871 | **0.0013** |

Data presented as mean ± SD.

App. = apparant; Disp. = displacement; Ult. = ultimate.

Two-way ANOVA: Bold values indicate significance with $p < 0.05$

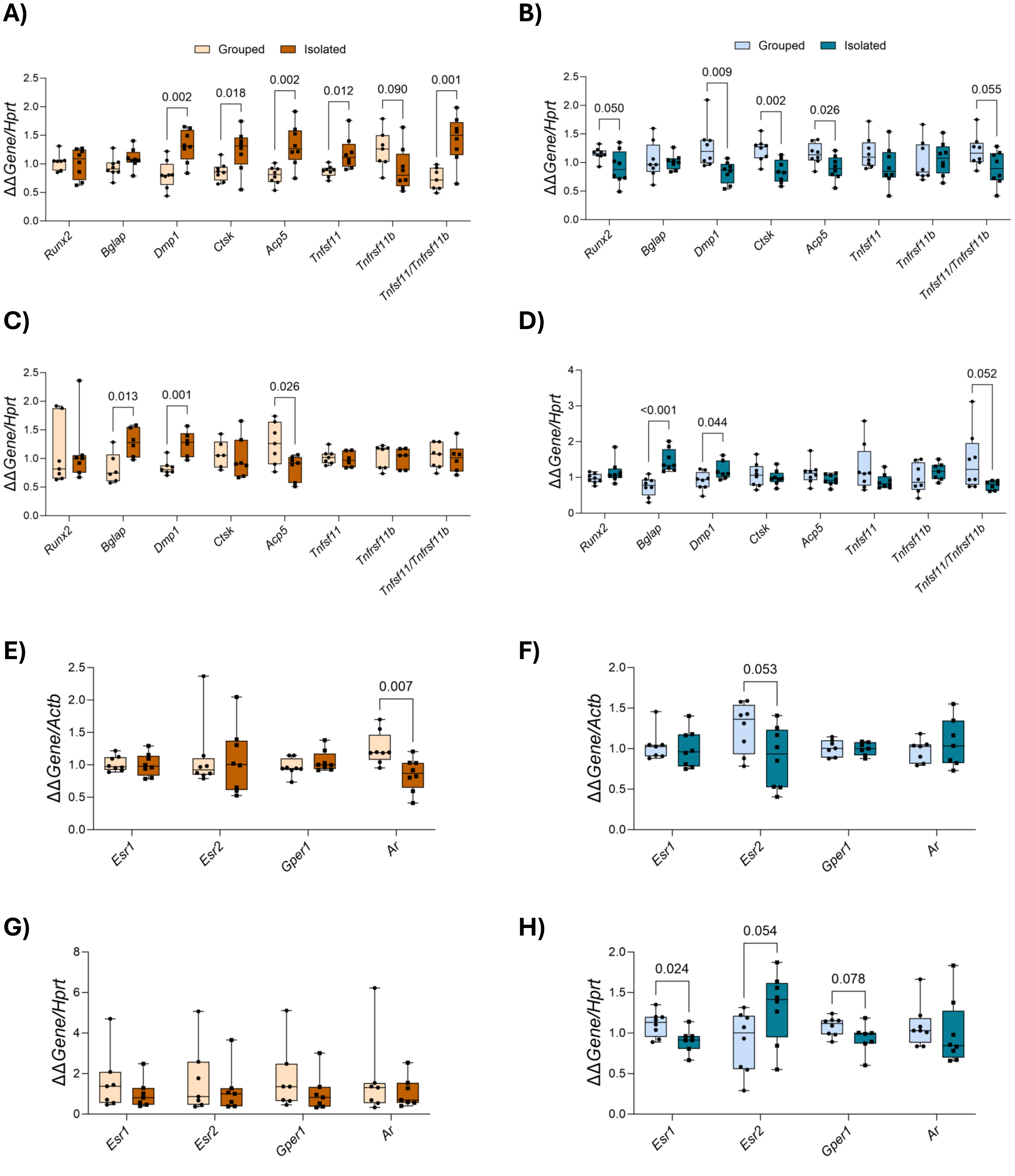


**Supplemental Figure 1. Supplemental bone turnover and sex hormone related gene expression**. Gene expression from whole tibia measured using RT-qPCR. Analyzed via multiple t-tests, comparisons with p ≤ 0.1 shown on graphs. N=6-8/group. **A-B)** 4-week female (A) and male (B) turnover-related gene expression. Normalized to *Hprt*. **C-D)** 8-week female (C) and male (D) turnover-related gene expression. Normalized to *Hprt*. **E-F)** 2-week female (E) and male (F) sex hormone-related gene expression. Normalized to *Actb*. **G-H)** 8-week female (G) and male (H) sex hormone-related gene expression. Normalized to *Hprt*.
